# Entanglement-based continuum conformational landscape of proteins

**DOI:** 10.64898/2026.08.17.745258

**Authors:** Paulina Malatesta, Roshita S. Chandnani, Jason Yalim, S. Banu Ozkan, Eleni Panagiotou

## Abstract

**Motivation:** With the rapid development of AI methods that predict protein structures from sequence, understanding the structure-function relation increasingly depends on quantitative structural descriptors that are both biologically meaningful and scalable to large datasets. Here, we introduce mathematical topology metrics that quantify the entanglement complexity of a tertiary protein structure while respecting uncrossability constraints.

**Results:** By employing only three such metrics across all protein structures in the Protein Data Bank, we represent the proteome structural space in a continuous three-dimensional space. Distances within this space capture structural similarity and correlate with functional similarity. We find that the mathematical entanglement based landscape of protein structural space diversifies with the evolutionary expansion of protein function across species. Moreover, this continuous representation reproduces CATH classifications with high accuracy for major structural classes. These results indicate that these metrics efficiently encode structural features linked to protein function and provide a more informative description than conventional metrics.

**Availability:** Data used in this study are available in the Protein Data Bank. Details of the machine learning model used can be found in https://github.com/roshitac/CATH_Classification-.

**Supplementary information:** Supplementary data are available at *Journal Name* online.

## Introduction

One of the main challenges in molecular biology is understanding the complex relationship between protein sequence, structure, and function [1, 2, 3, 4, 5]. Major advances in experimental methods have enabled the structure characterization of thousands of proteins, which are deposited in the Protein Data Bank (PDB) [6], while recent advances in machine learning enable the prediction of protein structure from sequence for millions of proteins [7]. Despite the vast amount of available data, connecting sequence variation to the fine-tuning of protein function in a predictive and interpretable manner remains a fundamental challenge. [8, 9]. The main challenge of computational protein design is to develop reliable models for protein generation and optimization that are based on computable features, something that has been explored to understand evolutionary relationships since the 1970s [10]. However, extracting biologically meaningful information from protein structure is substantially more difficult than from sequence alone [11]. Traditional approaches for comparing protein structures have primarily relied on amino acid sequence, Cartesian coordinates, dihedral angles, or simplified linear representations based on secondary structure elements. Yet, proteins are not static entities, but dynamic molecules that adopt many conformations [12, 13, 14, 15, 16, 17, 18, 19]. It is, therefore, challenging to determine whether structural differences reflect evolutionary divergence, functional difference or “experimental noise” [20, 21]. In this manuscript we employ relatively unexplored methods from mathematical topology to introduce a quantitative framework for characterizing protein structural complexity that captures information relevant to protein function.

The 3D tertiary structure topology of proteins is typically a non-mathematical descriptive framework, which focuses on understanding connectivity and arrangement of secondary structural elements within a protein. Mapping of these secondary structural elements and their relationships provides a reductionist view of complex 3D structures of proteins, and represents a powerful strategy for identifying recurring motifs, spatial arrangements and functional regions within proteins from different organisms and/or protein families [22]. From a mathematical perspective, methods from mathematical topology and geometry that represent proteins by point clouds have been used in studying proteins successfully [23, 24, 25, 26, 27, 28, 29]. These methods can derive many interesting results regarding shapes, such as voids, in the point cloud that represents a protein and connect those to effects of mutations and ligand-binding properties. In another mathematical topology approach, proteins can be represented by their C*α* atoms to give open piecewise linear curves in 3-space. This enables the application of methods from knot theory to characterize the complexity of proteins [30, 31, 32, 33, 34, 35, 36, 37, 38, 39, 40, 41, 42, 43, 44, 45]. By employing a closure approximation scheme it was found that almost 1% of the proteins in the PDB are knotted [46, 47], while by using closure approximation and scanning it was found that up to 9% of proteins have interesting topological features such as lassos, links, etc [35, 48, 49].

In this work, we employ novel methods from knot theory that enable us to quantify the topological complexity of proteins in a way that is applicable to all proteins, “knotted” or not. This complementary approach to previous studies focuses on representing proteins and their conformational ensembles by the point of view of a chain that satisfies uncrossability constraints and can adopt different conformations in a continuum. We use the Writhe, the second and the third Vassiliev measures, all of which are real valued metrics, to analyze the global mathematical geometry and topology of all proteins in the Protein Data Bank to provide a novel perspective to the structure-function relationship of proteins. The Writhe (similarly, the Gauss linking integral) has been applied successfully to proteins [50, 40, 51, 41, 42, 52, 53]. However, it is highly influenced by local geometrical characteristics of structure. Vassiliev measures capture the global conformational complexity for both closed and open curves in 3-space, and are continuous functions in the space of configurations (see [54]). Vassiliev measures of protein structure have been associated with secondary structure element packing [55] and protein kinetics [43]. These metrics enable to represent the protein tertiary structure topological and geometrical complexity in a 3-dimensional mathematical space, which we call the entanglement-based continuum conformational landscape of proteins. Using this framework, we find that proteomes across species evolve toward greater topological diversity. Furthermore, by integrating Gene Ontology (GO) functional annotations, we show that distances between proteins in this topological landscape correlate with functional similarity. By comparing our approach with the CATH database classification scheme, we show that CATH protein classes can be predicted with high accuracy using three entanglement based metrics and length alone. Together, these results establish continuum-based topological analysis of protein structure as not only mathematically rigorous, but also biologically meaningful. More broadly, this framework provides a quantitative link between protein structure and function and offers a scalable strategy for extracting biologically relevant insights from large protein-structure databases.

## Methods

### Entanglement-based metrics of protein structure complexity

Proteins are seen as piecewise linear curves with vertices their CA atoms. To maintain information only about structure, we analyze proteins, without any information regarding whether they are part of a multimeric complex or not. Also, we analyze proteins as a whole and not their domains. Protein backbone chains were assumed to be connected with a linear segment in-between gaps. Large gaps that affected the topology by introducing high *v*_2_, *v*_3_ values, were identified and the proteins excluded from the set. For each PDB file, we analyzed the first model, first chain. Of the 220 762 experimental protein structures that were downloaded from the PDB on June 7, 2024, a total of 211 829 matched our analysis criterion.

We employ three topological metrics to analyze protein structure complexity; the Writhe, the second Vassiliev measure and the third Vassiliev measure [56, 54, 43].

The Writhe of a curve in 3-space is defined as the Gauss linking integral over the curve [57]:

#### Definition 1

For an oriented curve. *ℓ* with arc-length parametrization *γ*(*t*), the Writhe, *Wr*, is the double integral over *l*:

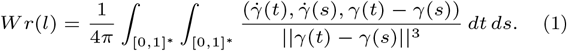

where integration is over all *s, t ∈* [0, 1], *s* ≠ *t*.

The Writhe is a real number, equal to the average algebraic sum of crossings in a projection of the chain (see Figure 1) over all possible projection directions. It can be computed exactly, avoiding numerical integration, using the algorithm described in [58]. The Writhe measures how much the chain turns around itself and it is a continuous function of the chain coordinates (not a topological invariant) for both closed and open curves.

**Fig. 1.**
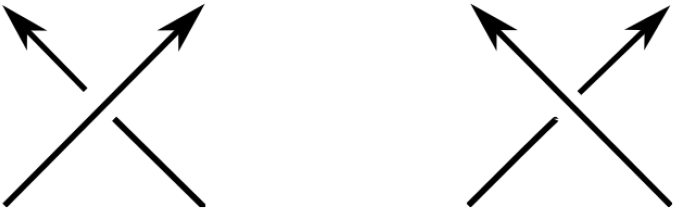
Crossing patterns. A projection of an oriented curve (protein backbone) results in crossings that can be assigned a positive (Left) or negative (Right) sign, depending on the orientation.

The second Vassiliev measure of open curves in 3-space is defined as follows [54]:

#### Definition 2

Let *l* denote a curve in 3-space with parametrization *γ*. The second Vassiliev measure of *l* can be expressed as:

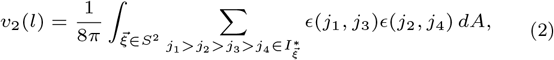

where *ϵ* (*s, t*) = *±*1, is the sign of the crossing between the projection of *γ*(*s*) and *γ*(*t*), and where 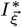 denotes the set of 4-tuples of alternating crossings (i.e., crossings such that if the projection of *γ*(*j*_1_) is over, resp. under, that of *γ*(*j*_3_), then the projection of *γ*(*j*_2_) is under, resp. over, that of *γ*(*j*_4_)) in the projection to the plane with normal vector 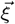.

The second Vassiliev measure captures a global pattern of crossings in a projection of a curve, averaged over all projections. For closed curves, the second Vassiliev measure is an integer topological invariant that can distinguish several knot types. For proteins, the second Vassiliev measure is a real number that is a continuous function of the chain coordinates in 3-space. As the endpoints of a curve tend to coincide, it tends to the value of the second Vassiliev invariant of the corresponding knot.

The third Vassiliev measure is defined as follows [54]:

#### Definition 3

Let *l* denote a curve in 3-space with. The third Vassiliev measure of *l* can be expressed as:

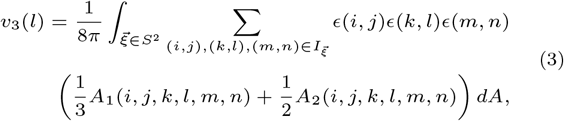

where 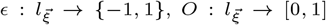 compute the sign and overstrand of a double point, respectively.

Where

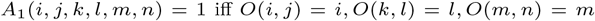

and

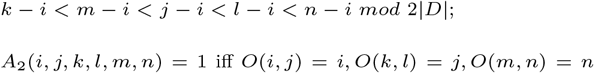

and

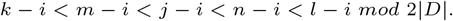

The third Vassiliev measure is a metric of structural complexity that can capture handedness. For example, in the case of knots, it can distinguish mirror images by sign. Together, the second and third Vassiliev measure can capture more refined topological characteristics of curves. Similarly to *v*_2_, the third Vassiliev measure is an invariant for knots that takes integer values. Together, *v*_2_ and *v*_3_ can distinguish many knot types and in particular, all of those found via the approximation closure scheme in the PDB [36]. For open curves, *v*_3_ is also a real number that is a continuous function of the curve coordinates. As the endpoints of a curve tend to coincide, *v*_3_ tends to the Vassiliev invariant of the corresponding knot. For *v*_2_ and *v*_3_, 5000 random projections were used for their computation.

Examples of *Wr, v*_2_, and *v*_3_ values for specific proteins are shown in Figure 2. All three proteins have approximately the same length but differ in their Writhe and Vassiliev measure values. Their Vassiliev measures indicate that they represent examples of increasing complexity. The first two proteins (7tx2, 8ae9) have similar Writhe values but 8ae9 has a more complex global structure than 7tx2, as indicated by their Vassiliev measures. The last protein, 6r6y, has lower Writhe value, indicative of smaller local geometrical complexity, but higher *v*2, *v*3 values, which indicate knottedness of the trefoil type. Note that the first two proteins are considered trivial in the knot closure approach, while the last protein is considered a trefoil knot.

**Fig. 2.**
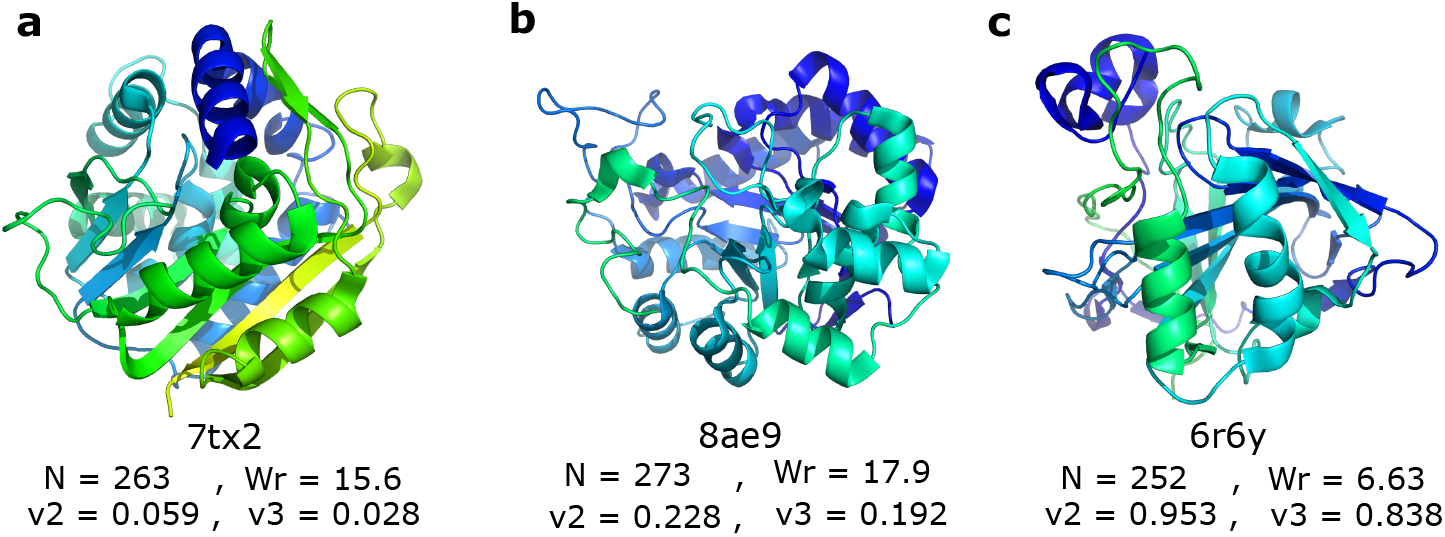
Proteins with their corresponding PDB IDs, length (N), Writhe (W r), second, v2, and third, v3, Vassiliev measure values. All these proteins share a similar length but have distinct topological and geometrical characteristics, as depicted by their W r, v2, v3 values. High W r values indicate mostly higher local geometrical complexity, with α-helices contributing to higher W r values. Higher v2, v3 values indicate higher global complexity and even knotting, as in the case (c). Note that the right handed trefoil knot has v2 = v3 = 1.

### Distance-based metrics of structural similarity of proteins

RMSD and TM-score are quantitative metrics of structural similarity between two proteins. Typically, as a first step the proteins are superimposed [59].

#### Definition 4

Root mean square deviation (RMSD) is defined as:

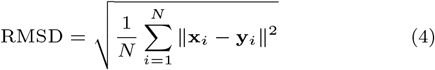

where *N* is the number of matched atoms, **x**_*i*_ and **y**_*i*_ are the coordinate vectors of the *i*-th matched atoms in the first and second structures, respectively, and ‖. ‖ denotes the Euclidean norm.

#### Definition 5

Template Modeling score (TM-score) is defined as:

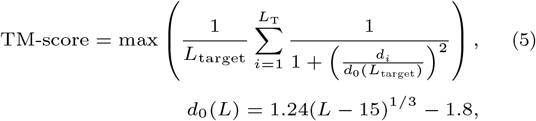

where *L*_target_ is the target length, *L*_T_ is the number of aligned residues to the template structure, *d*_*i*_ is the distance between the *i*-th pair of aligned residues, and *d*_0_ is a scale that normalizes distances.

### Lin measure of protein function similarity

Protein function similarity between two proteins can be quantified by comparing their Gene Ontology terms via the Lin measure [60]:

#### Definition 6

Lin measure of function similarity of proteins is defined as as:

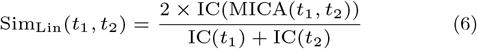

where

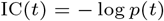

is the information content, and:

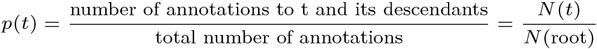

where *N* (*t*) denotes the number of proteins annotated with the term *t* or any of its descendants, and where MICA is the most informative common ancestor of the two compared GO terms that has the maximal information content.

Higher IC(MICA) refers to more specific common function (less populated in the genome). GO terms that are higher up (closer to the root) in the GO graph (DAG) should have low IC.

## Results

We present data on the mathematical analysis of protein structures deposited in the Protein Data Bank using three mathematical topology measures (the Writhe, the second and third Vassiliev measures), that capture the complexity of a protein structure seen as a string of C*α* atoms. A total of 211 829 protein structures were analyzed (see Methods Section).

### The continuum entanglement-based landscape of proteins

The entanglement-based landscape of proteins is shown from one perspective in Figure 3 (see also Section 1 in the SI). Each protein in the PDB can be represented by a point in a 3-dimensional topological space of axes *Wr/N, v*_2_, *v*_3_. (A normalization of *Wr* by *N* is used to account for the high dependency of *Wr* on *N*). Similarly, we can use a representation in a 4-dimensional space, *Wr, v*_2_, *v*_3_, *N*. Of the 211 829 proteins analyzed from the PDB, a total of 6569 proteins (3.1%) had a value of *v*_2_ and *v*_3_ equal to zero, while no protein had a Writhe value equal to zero. Thus, all proteins have a non trivial geometric complexity (the latter is reflected by the Writhe axis, see Figure 1 in the SI) and 96.4% of proteins have a non-trivial topological entanglement global structure (reflected by the *v*_2_, *v*_3_ plane). Proteins of low global entanglement complexity (yet non-zero) are observed near the origin in the *v*_2_, *v*_3_ plane and those with higher topological complexity, are observed away from the origin. An area [−1, 1] *×* [−1, 1] centered at the origin of the projection of the entanglement landscape in the *v*_2_, *v*_3_ plane is shown in Figure 3. The general observed pattern of the *v*_2_, *v*_3_ plane resembles that of Willerton’s fish [61], due to the underlying correlations of the second and third Vasisliev measures (see Section 2 in the SI for more information). We find that 98.9% of analyzed proteins have a value in a disk of radius 0.5 from the origin in the (*v*_2_, *v*_3_) plane, while 83.5% have values of *v*_2_ and *v*_3_ within a ball of radius 0.1. 55.5% of all proteins have a positive *v*_3_ value, indicating a small overall preference for what could be described as right-handed global conformations. This tendency increases when we restrict proteins to 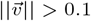, where 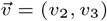. Of the 34 890 proteins with *IvI >* 0.1, 21 586 (62%) had *v*_3_ *>* 0, These results suggest that higher global topological complexity may be associated with an increased bias towards right handed global conformations. Finally, proteins that contain knots are found at the neighborhood of points in ℤ *×* ℤ *\* (0, 0) in the *v*_2_, *v*_3_ plane.

**Fig. 3.**
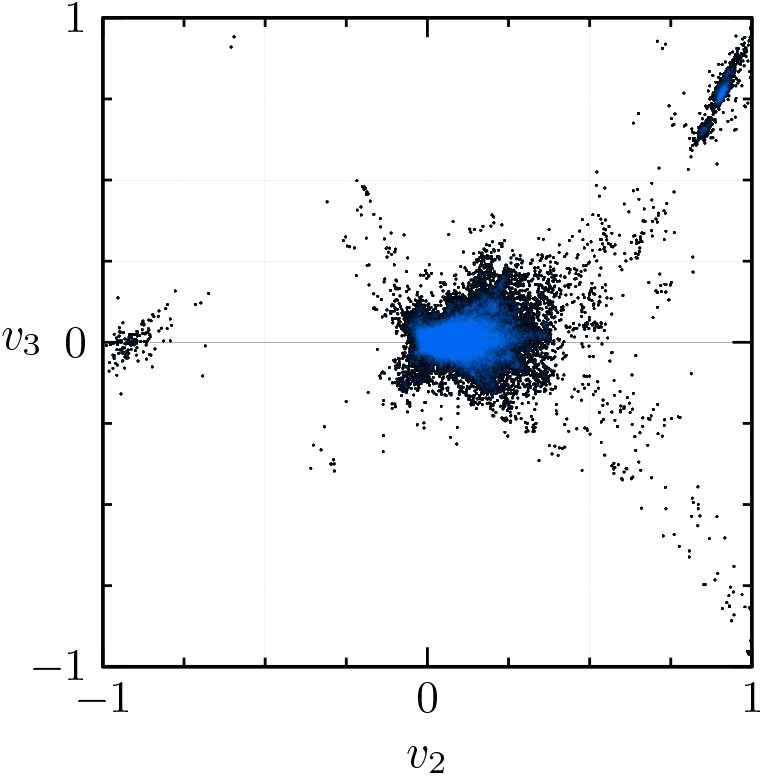
Projection of the entanglement-based landscape of proteins in the PDB on the v2, v3 plane. The heatmap indicates with blue the regions of higher density. The general observed arm-like structure represents the underlying correlation between v2, v3, but asymmetries around the axes and discontinuous island-like structures represent PDB protein-specific patterns. See also Section 1 in the SI.

#### Euclidean distance in the topological landscape captures protein structure similarity

In this section we compare the quantitative topological similarity of proteins to the Root Mean Square Deviation (RMSD) [62, 63] and to the Template Modeling score (TM-score) [64, 59] (see Methods Section for definitions). We stress that both RMSD and TM-score require an alignment of the structures prior to computing [62, 59].

Let *prot*_1_, *prot*_2_ denote two proteins which correspond to the points 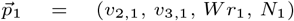 and 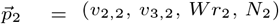 in the continuum mathematical topological landscape, respectively. The coordinates of the 4D vectors are the second and third Vassiliev measures, *v*_2_, *v*_3_, the Writhe (*Wr*) and length (*N*), respectively. We define the topological distance of two proteins in the continuum mathematical topological landscape to be the standardized Euclidean distance 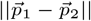, namely:

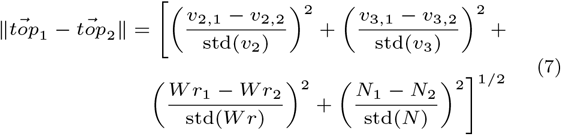

where *std* is the standard deviation of *v*_2_, *v*_3_, Writhe and length of the distribution of metrics in the entanglement-based landscape. When focusing on a subset of the topological landscape, such as proteins within one specific family, we define the topological distance as the standardized Euclidean distance with respect to the particular subset.

To compare the performance of RMSD and TM-score with that of topological distance in assessing protein structure similarity, we focus on one of the largest families in the PDB, Aldolase, for which we compute the the RMSD, TM-score and topological distance of every pair of proteins therein. The results are shown in Figure 4. We find that even small RMSD values, of less than 5, may correspond to a large topological distance of protein structures. This indicates that structures that are considered similar by RMSD metrics, may still have quantitatively distinct entanglement topologies. Similarly, proteins with high TM-score, over 0.5, which are otherwise considered to have the same fold, can have large quantitative differences in topological structures that can be detected with the mathematical metrics proposed here. This method of comparing protein structures is thus not only faster than other methods, but also likely more accurate and could be used as a stand-alone metric or in combination with AI methods of comparison of structures [65].

**Fig. 4.**
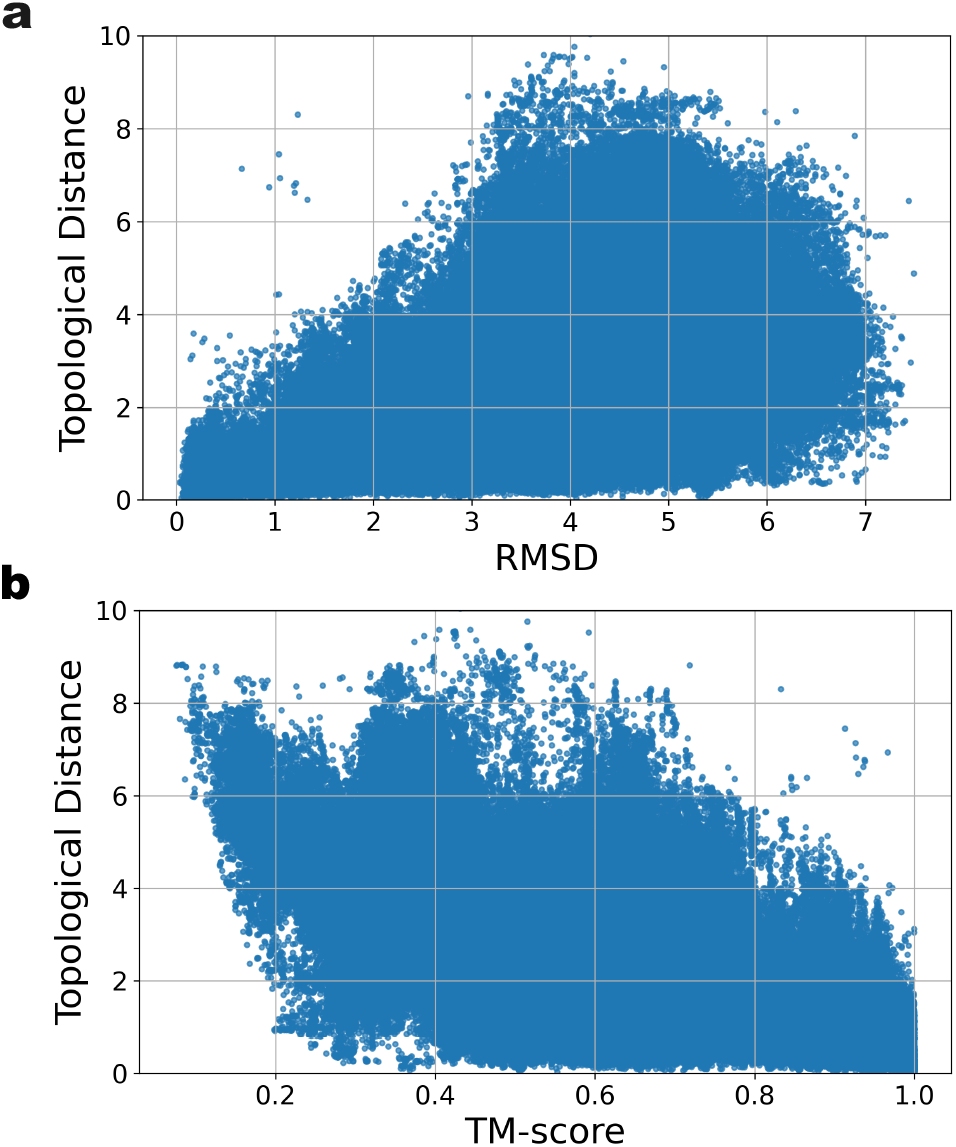
Comparison between RMSD and TM-score with Topological Distance. (a) RMSD and (b) TM-score versus Topological Euclidean distance in the entanglement-based protein landscape for all protein pairs of the Aldolase family. We observe that proteins with low RMSD and high TM-score, which are otherwise considered to have the same fold, can have large entanglement-based structural differences.

### Protein evolution and function in the topological landscape

#### Protein evolution in the continuum mathematical topological landscape of proteins

It is known that, across different species, the distribution of protein amino acid chain lengths varies relatively little, particularly when considered a distinguishing characteristic among organisms. Since the (global) Writhe of a protein is highly sensitive to chain length, it is difficult to decouple length effects from structural complexity using it. In contrast, the Vassiliev measures *v*_2_ and *v*_3_ capture global protein structural complexity that is less directly dependent on length, making them suitable quantities for examining the role of structural complexity in protein evolution.

To investigate how the quantitative diversity of structures within the topological landscape correlates with evolution, we analyzed proteins from three species that are both extensively represented in the PDB and broadly reflect increasing biological and functional complexity: *E. coli*, yeast, and humans. *E. coli* represents a prokaryotic organism with the core biochemical machinery required for cellular life. Yeast serves as a unicellular eukaryotic model organism, exhibiting greater organizational and functional complexity than bacteria while remaining simpler than multicellular systems. Humans represent highly complex multicellular eukaryotes with extensive cellular specialization and regulatory diversity. Human proteins constitute a substantial fraction of structures deposited in the PDB [66], reflecting the longstanding emphasis on human health, disease mechanisms, and drug discovery [67]. Yeast has become a central model system for eukaryotic biology because of the well-established relationships between its genes, proteins, and cellular functions [68]. Similarly, *E. coli* remains one of the most extensively studied organisms due to its versatility, rapid growth, and experimental accessibility [69].

Figure 5 shows the Cumulative Distribution Function (CDF) for *v*_2_ and *v*_3_ for values in the ranges [-0.1, 0.4] and [-0.3, 0.3], respectively, as well as their corresponding histograms. *v*_2_ values were fitted to a skewed-normal distribution, while *v*_3_ to a normal distribution. Note that the skewness of *v*_2_ is inherent to the mathematical properties of the second Vassiliev invariant and is also observed for random walk systems (see discussion in SI). The results show that the variance of *v*_2_ and *v*_3_ differs by organism. Specifically, we notice that human has the largest variance while *E. coli* the smallest. This progressive increase in variance from prokaryotes to multicellular eukaryotes suggests that protein evolution expands the accessible topological landscape of protein structures. In turn, this increasing topological diversity is consistent with the emergence of broader and more specialized functional repertoires during evolution.

**Fig. 5.**
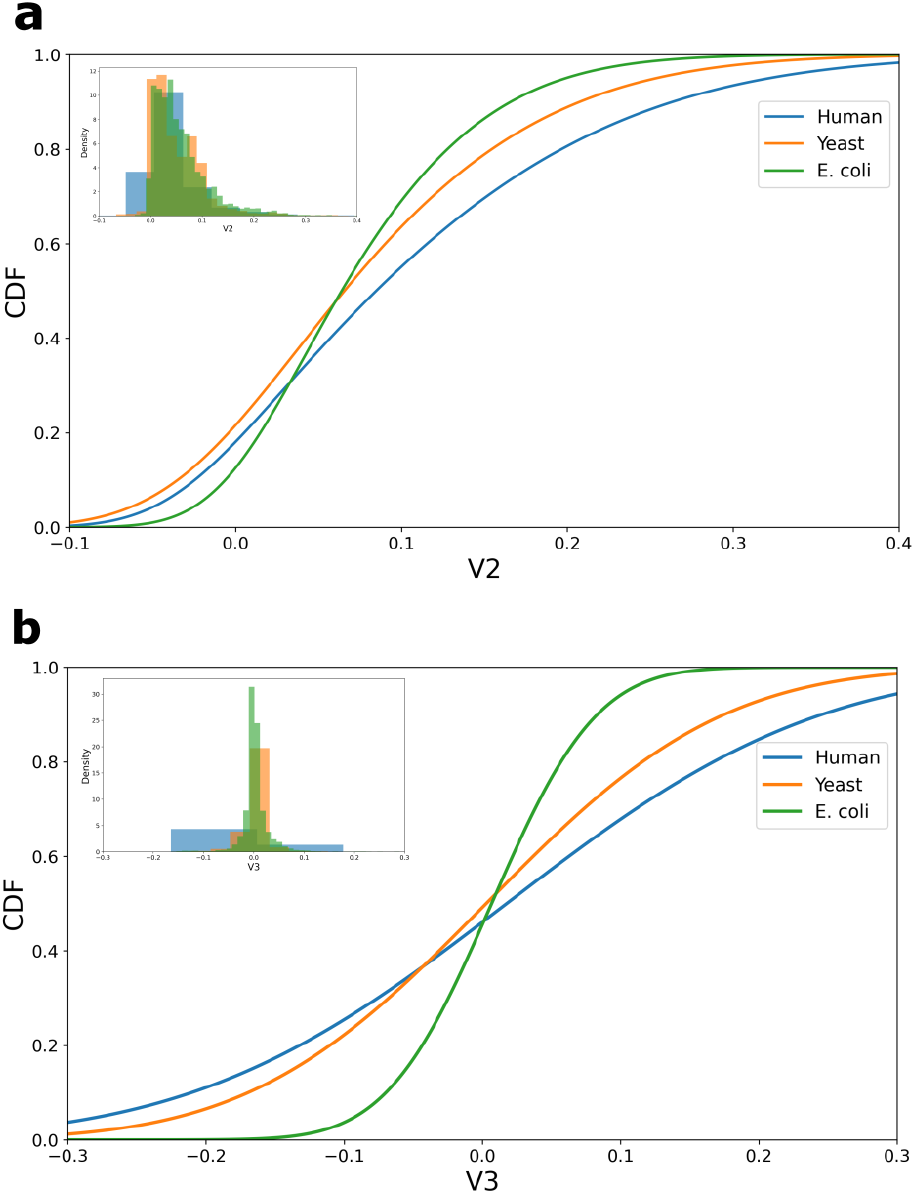
The distribution of entanglement-based metrics for proteins of different organisms. **(a)** Cumulative Distribution Function of v2 and (b) Cumulative Distribution Function of v3, along with their corresponding histograms for Human (Blue), Yeast (Orange), and E. coli (Green) proteins. The variance increases with evolution, suggesting that proteins adopt more diverse structures, as they acquire more diverse functions.

#### The entanglement-based landscape of proteins and protein function

An important point in searching for protein structure descriptors, is to find those that can compare proteins in a way that reflects their function similarity. We examine whether the distance of proteins in the quantitative topological landscape correlates with function similarity [60].

Protein function can be determined through Gene Ontology (GO) [70], which provides information about the molecular functions, cellular locations, and processes gene products may carry out.

A common method to assess protein function similarity is via the Lin measure [60, 71, 72] (see Methods Section for definition). The Lin measure of function similarity is defined using the GO annotations of proteins and takes values between 0 and 1. The higher the Lin score, the more functional similarity between two GO terms. It has been shown that most pairs of proteins exhibit lower scores (0.15) and an average score no higher than 0.4 [73]. Since proteins can have many GO annotations, the Lin similarity between two proteins can be calculated by computing the Lin similarity of all terms of one protein compared to all the terms of the other protein and taking the average. Since GO terms are assigned at the Uniprot sequence level and multiple PDB IDs may correspond to the same Uniprot ID, in this study, all PDBs that share the same Uniprot ID were treated as replicates as the same protein and an average of their topological values were taken.

We focus on four regions in the entanglement landscape of proteins that capture a wide range of distinct topological entanglement states. In particular, we focused on the regions: 0 *< v*_2_, *v*_3_ *<* 0.08 (Low Topological Complexity), 0.08 *< v*_2_, *v*_3_ *<* 0.5 (Intermediate Topological Complexity), 0.5 *< v*_2_, *v*_3_ *<* 0.9 (Transition to Knotting), 0.744 *< v*_2_ *<* 1.22 and 0.585 *< v*_3_ *<* 1.2 (Right Handed Trefoil Knot), while the Writhe and length, *N*, of proteins are not bounded. Knotted protein region bounds are defined so that they are in agreement with literature of knotted proteins [36]. The 0 *< v*_2_, *v*_3_ *<* 0.08 region contains 95.8% of the total proteins in the four regions, namely, 5927 unique Human Uniprot IDs (in total 41 710 Human proteins), 1342 unique Yeast Uniprot IDs (10 672 Yeast proteins in total), and 1005 unique *E. coli* Uniprot IDs (10 434 *E. coli* proteins in total). The 0.08 *< v*_2_, *v*_3_ *<* 0.5 region contains 122 Human, 33 Yeast and 34 *E. coli* unique Uniprot IDs (a total of 688, 57, and 160 proteins, respectively). The 0.5 *< v*_2_, *v*_3_ *<* 0.9 (Transition to Knotting) region contains 315 Human proteins (only 9 unique Uniprot ids). The Right-Handed Trefoil region contains 84 unique Human Uniprot IDs (a total of 1362 Human proteins), 8 unique Yeast Uniprot IDs (15 Yeast proteins in total), and 10 unique *E. coli* Uniprot IDs (22 *E. coli* proteins in total).

Figure 6 shows the relationship between topological distance and Lin functional similarity for protein structures from the main organisms studied (human, yeast, and *E. coli*) across three distinct regions of the protein topological landscape: Low Topological Complexity, Intermediate Topological Complexity, and the Right-Handed Trefoil region. Across all protein regions (namely, all entanglement complexity ranges), including those shown in the SI, we observe a consistent relationship between structural and functional similarity: higher values in topological distance (pairs of proteins with low structure entanglement similarity), correspond to lower Lin similarity values (low functional similarity). Similarly, smaller values in topological distance (pairs of proteins with high structure entanglement similarity), correspond to higher Lin similarity values (high functional similarity). In particular, Figure 6(a) shows the relation between Topological distance and function similarity for the low topological complexity region that contains a large number of proteins. Consistently with the other regions, we observe an increase in protein function similarity (high Lin similarity score) with decreasing topological distance of proteins. Particular to this dataset, which focuses on very low entanglement complexity structures, is that Lin similarity exceeds 0.5, which is the typical high similarity score in most datasets. This occurs for only 0.02% of the data in this region and may be due to the fact that many of the proteins in this low topological complexity set have only general function specifications. In all regions, we observe that the trend is more pronounced for Human proteins, possibly reflecting the larger sample size as well as their higher function diversity, which contributes to a better distribution of proteins in the Lin similarity ranges. Overall, these results establish that the proposed quantitative metrics of protein structural complexity are biologically meaningful and strongly associated with protein function. Importantly, this relationship holds across the full range of topological complexity examined, from minimally entangled structures to knotted proteins.

**Fig. 6.**
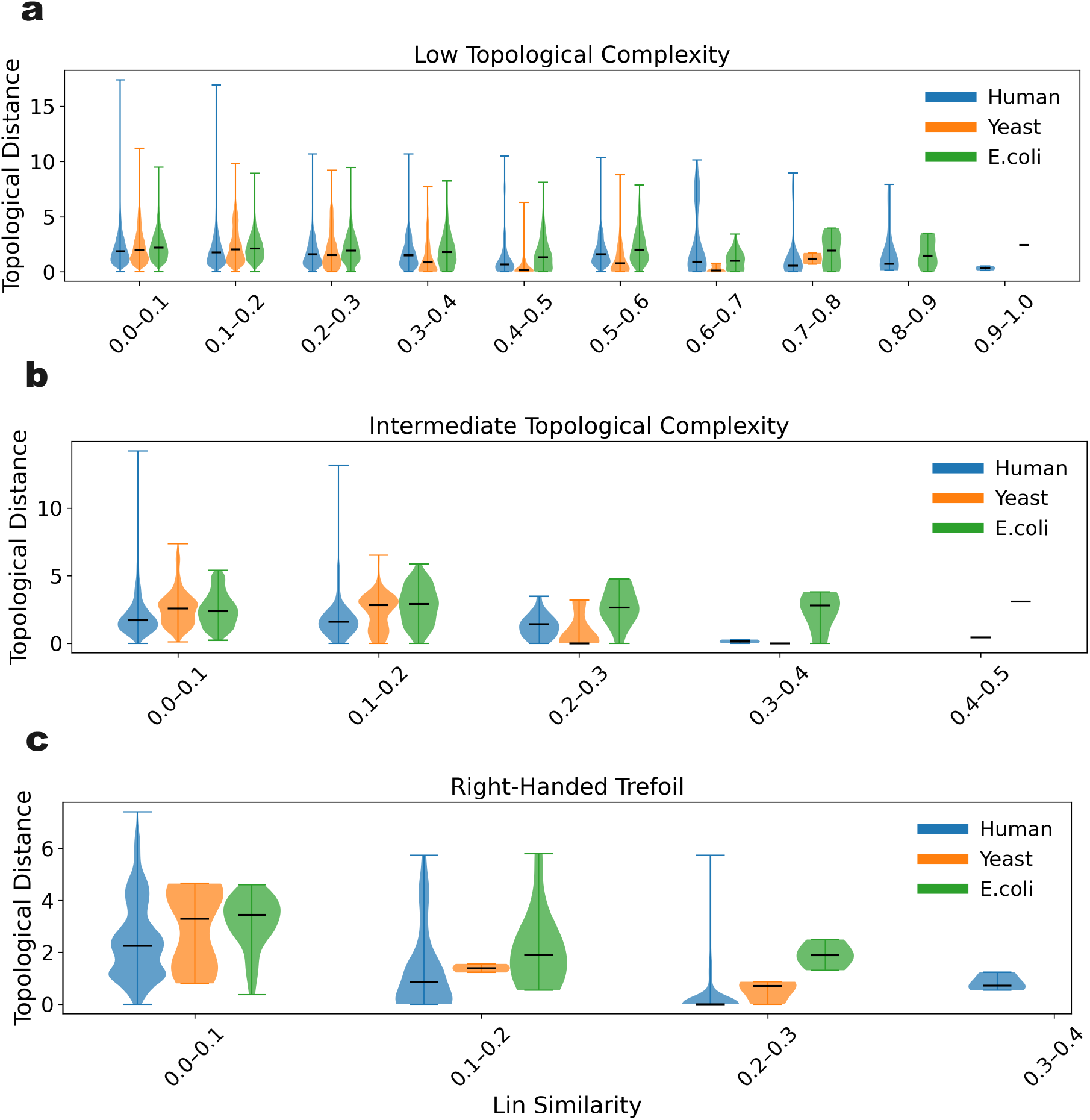
Topological distance of proteins versus Lin similarity. Euclidean distance of proteins in the topological landscape versus function similarity grouped by organism (Human, Yeast and E. coli) for low topological complexity ((a) 0 < v2, v3 < 0.08), intermediate topological complexity ((b) 0.08 < v2, v3 < 0.5), and “knotted” ((c) Right-Handed Trefoil) proteins. In all cases, we see that increasing protein function similarity correlates with decreasing distance in the topological entanglement landscape.

Figure 7 shows the topological distance of proteins as a function of their function similarity grouped by organism. All plots agree with the observations derived from Figure 6, namely, topological distance correlates with functional similarity. In particular for human and yeast proteins, we find that the larger the topological distance the lower their Lin similarity is. For Human proteins, we also include the region 0.5 *< v*_2_, *v*_3_ *<* 0.9, that captures a transition to higher topological complexity region, which also shows the same trend. For E.coli we see a similar trend, but it is not as evident in comparison to the other two organisms. This finding is consistent with the fact that E.coli is the simplest organism among the three, therefore it exhibits less complexity and less functional evolution and differentiation.

**Fig. 7.**
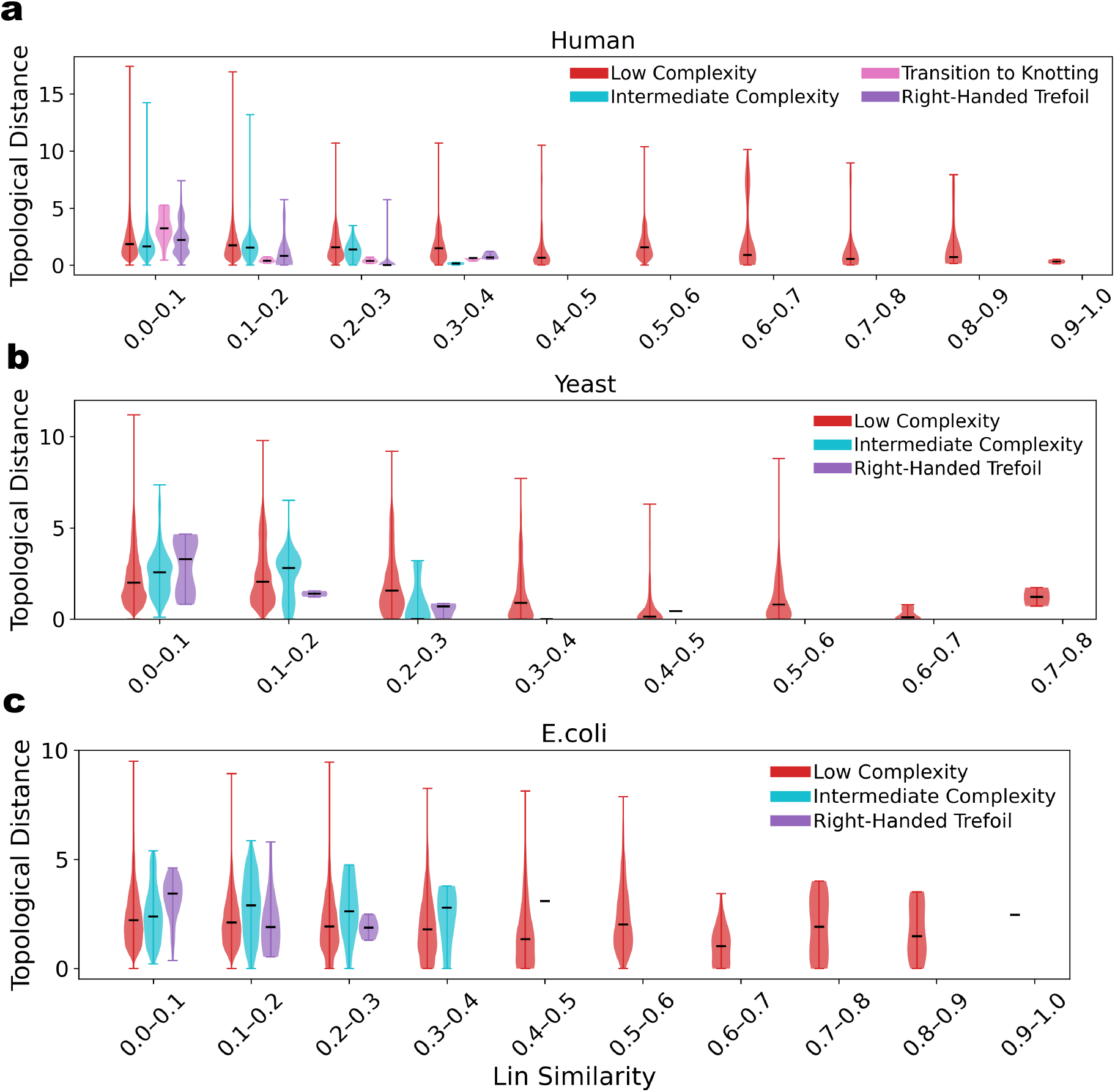
Topological distance of proteins versus Lin similarity grouped by organism. Euclidean distance of proteins in the entanglement based landscape versus function similarity for (a) Human, (b) Yeast, and (c) E. coli proteins in ranges of low, intermediate and high entanglement complexity. In all cases, we see that increasing protein function similarity correlates with decreasing distance in the entanglement landscape.

### CATH protein classifications and the topological landscape

Protein classification frameworks such as CATH have been widely used to infer structural, functional, and evolutionary relationships between proteins [74, 75]. These classifications typically combine secondary-structure composition, fold organization, sequence similarity, and structure-alignment methods [76]. In parallel, continuum geometric and topological descriptions of proteins have emerged as quantitative approaches for characterizing protein structural complexity [77]. Here, we investigate how global entanglement-based descriptors of protein structure relate to the hierarchical classifications of the CATH database.

A total of 127 907 structure annotations were downloaded via the PDBe CATH API. Of these, 123 379 were in the 211 829 proteins analyzed in this study. The CATH (Class, Architecture, Topology, Homology) hierarchy organizes proteins according to secondary-structure composition, overall structural arrangement, fold connectivity, and evolutionary relationships. To focus on single-domain structural organization, proteins containing multiple domains were removed from the analysis, reducing the dataset by: 28% for *α* proteins, 24% for *β* proteins, and 41% for *α*-*β* proteins.

We incorporate the Writhe (Wr), the second and third Vassiliev (*v*_2_, *v*_3_) measures and the length of proteins (N) as input features for classification. A multilayer perceptron (MLP) classifier was trained on standardized features using two hidden layers of 100 and 50 neurons with ReLU activation and a softmax output layer. The Adam optimizer (learning rate 10^−3^) and L2 regularization (*λ* = 0.001 or 0.01) were employed for up to 500 epochs, with hyperparameters selected through 3-fold cross-validation. The dataset was divided into training, validation, and test sets, and no evidence of overfitting was observed (see Section 3 in the SI).

Using only the four descriptors (*Wr, v*_2_, *v*_3_, *N*), the model predicts the three dominant CATH classes (corresponding to “C”) with a test accuracy of 0.892 *±* 0.002 (range 0.888–0.897), a macro-F1 score of 0.8916 *±* 0.002, and a weighted-F1 score of 0.8926 *±* 0.002 (see Figure 8). Classification performance further improves in pairwise comparisons between dominant classes. In particular, the model achieves accuracies of 92.7% for *α* versus *α*-*β*, 92.5% for *β* versus *α*-*β*, and 97% for *α* versus *β* classification. We next examined deeper levels of the CATH hierarchy for these dominant classes. The corresponding results are summarized in Table 1 and Figure 9 (see also SI). We note that for *α*-*β* proteins, class 3.40 accounts for approximately 45% of the dataset; we report results both including and excluding this dominant family.

**Fig. 8.**
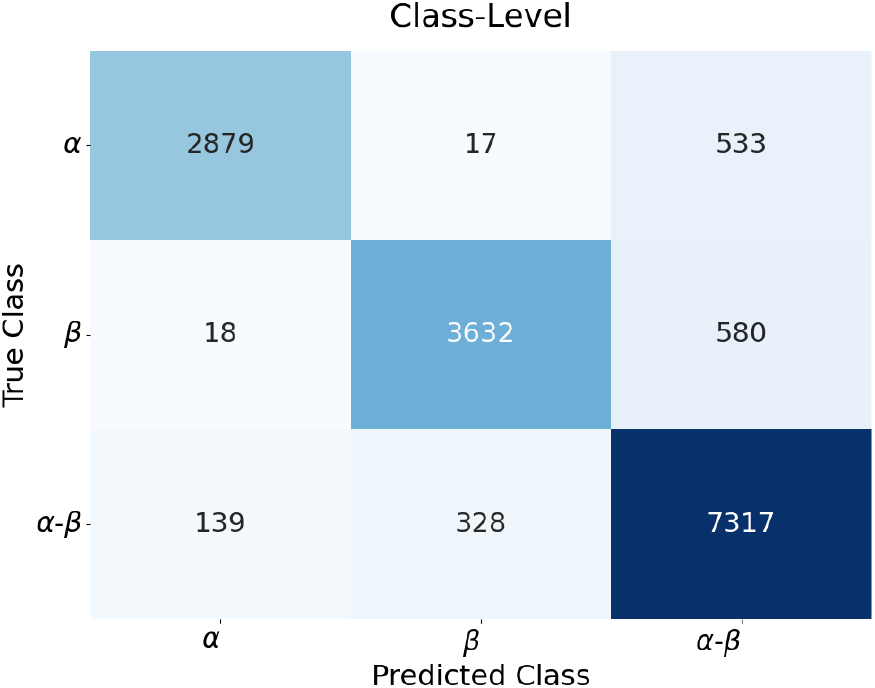
Confusion Matrix for Class-Level prediction of proteins, using entanglement based metrics. An ML model based on wr, v2, v3, N achieves high accuracy in predicting a protein’s class.

**Table 1.** MLP classification performance across the CATH hierarchy for three classes (*α, β, α-β*). All metrics are reported with their mean *±* one standard deviation over thirty stratified eighty–twenty train-to-test splits. The absolute difference between training and test accuracy was less than 3% across all settings.

| Protein structural domain | Level | # Classes | # Proteins | Test Accuracy | Acc. Min–Max | Macro-F1 | Weighted-F1 |
| --- | --- | --- | --- | --- | --- | --- | --- |
| $\alpha$ | C.A | 3 | 16 926 | $0.856 \pm 0.006$ | 0.841–0.868 | $0.778 \pm 0.011$ | $0.853 \pm 0.006$ |
| | C.A.T | 13 | 10 567 | $0.841 \pm 0.007$ | 0.821–0.860 | $0.802 \pm 0.009$ | $0.837 \pm 0.007$ |
| | C.A.T.H | 10 | 8029 | $0.917 \pm 0.005$ | 0.906–0.926 | $0.901 \pm 0.006$ | $0.917 \pm 0.005$ |
| $\beta$ | C.A | 6 | 19 411 | $0.785 \pm 0.007$ | 0.773–0.800 | $0.709 \pm 0.012$ | $0.781 \pm 0.007$ |
| | C.A.T | 8 | 15 212 | $0.817 \pm 0.007$ | 0.803–0.833 | $0.803 \pm 0.009$ | $0.816 \pm 0.007$ |
| | C.A.T.H | 7 | 9226 | $0.918 \pm 0.007$ | 0.904–0.937 | $0.901 \pm 0.009$ | $0.918 \pm 0.007$ |
| α-β | C.A | 7 | 38 447 | $0.756 \pm 0.006$ | 0.741–0.768 | $0.649 \pm 0.015$ | $0.751 \pm 0.007$ |
| | C.A.T | 13 | 23 308 | $0.763 \pm 0.006$ | 0.752–0.778 | $0.656 \pm 0.010$ | $0.756 \pm 0.006$ |
| | C.A.T.H | 12 | 12 554 | $0.781 \pm 0.007$ | 0.766–0.796 | $0.781 \pm 0.008$ | $0.780 \pm 0.007$ |
| α-β 3.40 | C.A.T | 5 | 13 702 | $0.903 \pm 0.005$ | 0.893–0.913 | $0.815 \pm 0.011$ | $0.901 \pm 0.005$ |
| | C.A.T.H | 8 | 8644 | $0.809 \pm 0.011$ | 0.779–0.834 | $0.800 \pm 0.012$ | $0.808 \pm 0.011$ |
| α-β (3.40 Excluded) | C.A.T | 11 | 10 940 | $0.836 \pm 0.008$ | 0.822–0.856 | $0.7635 \pm 0.013$ | $0.835 \pm 0.008$ |
| | C.A.T.H | 9 | 6081 | $0.894 \pm 0.009$ | 0.875–0.912 | $0.885 \pm 0.001$ | $0.893 \pm 0.009$ |

**Fig. 9.**
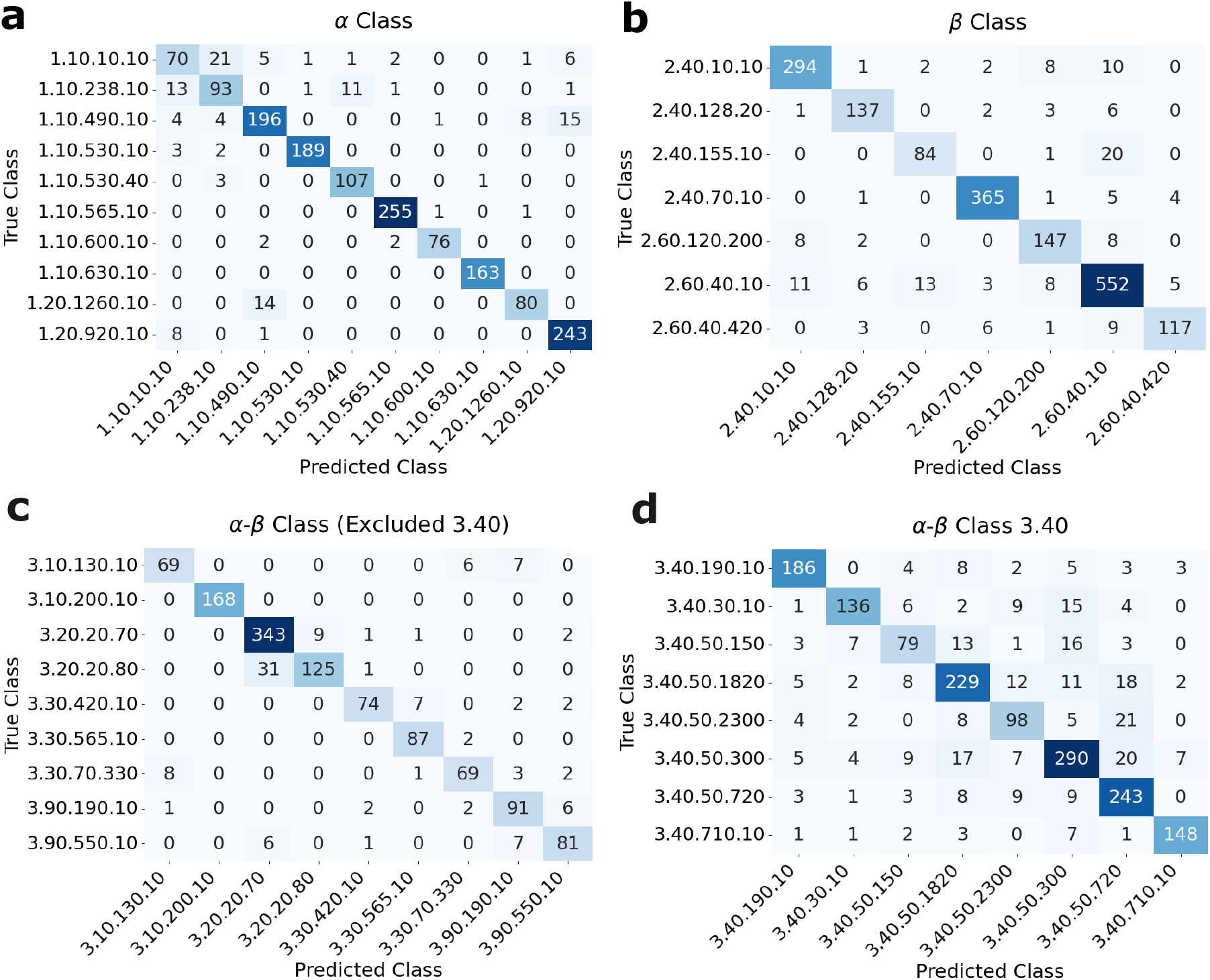
Confusion matrices of CATH protein classification prediction based on entanglement metrics for the three main classes. (a) *α* class (b) *β* class (c) *α*-*β* proteins excluding class 3.40, which consists of 44.9% of the data (d) *α*-*β* class 3.40.

To understand the relative contribution of each descriptor, features were sequentially added to the model. Across all cases, the standard deviation of cross-validation scores remained below 1%, indicating stable model performance. For *α* proteins, Writhe and protein length account for most of the predictive performance, while inclusion of *v*_2_ and *v*_3_ substantially improves CATH Topology classification from 76.9% to 83.6%. Similarly, for *β* proteins, Writhe and length dominate Class-level discrimination, whereas the addition of *v*_2_ and *v*_3_ improves Topology and Homology classification from 84.3% to 91%. The strongest effect is observed for *α*-*β* proteins, where Writhe and length alone provide only partial discrimination. Incorporating *v*_2_ and *v*_3_ increases Architecture classification accuracy from 68% to 75.9%, Topology classification from 67.1% to 78.4%, and Homology classification from 65.7% to 78%. These improvements are even larger when excluding the dominant 3.40 family. The stronger dependence on higher-order entanglement descriptors in *β* and *α*-*β* proteins may reflect the greater structural complexity and nonlocal connectivity characteristics of these architectures relative to predominantly *α*-helical proteins.

Overall, these results demonstrate that low-dimensional continuum entanglement descriptors encode substantial information about protein structural organization across multiple levels of the CATH hierarchy. While class-level discrimination is driven primarily by Writhe and protein length, higher-order Vassiliev measures become increasingly important for resolving Topology and Homology relationships, particularly in *β* and *α*-*β* proteins. Importantly, these classifications are obtained without sequence information, structural alignment, or feature engineering based on protein interactions and structural annotations, relying solely on the geometry of the C*α* backbone chain. These findings demonstrate that continuum topological descriptors not only capture biologically meaningful aspects of protein architecture, but also provide a rigorous, quantitative, and computationally efficient framework for protein classification that complements existing AI-based approaches employing extensive engineered structural and interaction features [78, 79, 80, 81, 82, 83].

## Discussion

We introduced a continuum topological entanglement framework for quantifying protein structural complexity by modeling proteins as linear open curves in three-dimensional space. Using only three global entanglement descriptors, the Writhe and the second and third Vassiliev measures, together with protein length, we constructed a low-dimensional representation of the structural organization of proteins across the Protein Data Bank. Unlike conventional approaches that rely on sequence similarity, structural alignment, or engineered structural descriptors, this framework characterizes proteins directly from the geometry of the C*α* backbone chain. Moreover, these metrics capture a protein’s connectivity and uncrossability constraint, reflecting aspects of its entropy.

Our analysis shows that proteins occupying nearby regions in the entanglement landscape tend to exhibit similar structural and functional properties. In particular, we find that topological distance correlates with functional similarity measured through Lin similarity, suggesting that proteins with related functions also share similar global entanglement organization. Moreover, comparisons with RMSD- and TM-score-based analyses indicate that the continuum entanglement representation captures meaningful aspects of structural similarity while remaining independent of alignment procedures. We further observe that the diversity of the topological descriptors increases from *E. coli* to Yeast to Human proteins, consistent with the idea that increased structural and functional diversification during evolution is accompanied by a broader exploration of protein entanglement space. The continuum entanglement descriptors also recover substantial information across multiple levels of the CATH hierarchy. Using only Writhe, *v*_2_, *v*_3_, and protein length, a simple neural-network classifier achieves strong performance in distinguishing major protein classes and deeper CATH subclassifications, suggesting that global entanglement descriptors encode biologically relevant aspects of protein architecture beyond secondary-structure composition alone. Overall, these findings demonstrate that low-dimensional continuum topological descriptors capture biologically meaningful aspects of protein organization, classification, and evolution within a rigorous and computationally efficient framework. Because these descriptors are continuous in conformational space, they naturally extend to protein dynamics and conformational ensembles, providing a complementary perspective on the sequence–structure–function relationship in proteins.

