## Supplementary Information for "Entanglement-based continuum conformational landscape of proteins"

August 17, 2026

### 1 3D visualization of the entanglement-based conformational landscape of proteins

The conformational landscape of proteins in the PDB in $Wr/N, v_2, v_3$ is shown in Figure 1 where the colors correspond to distinct species. The interactive figure is available in Entanglement landscape by Species plot. An interactive figure with the topological landscape colored by protein main Class $(\alpha, \beta, \alpha\beta)$ are shown in Entanglement landscape by Class plot and by CATH classification in Entanglement landscape by CATH classification plot.

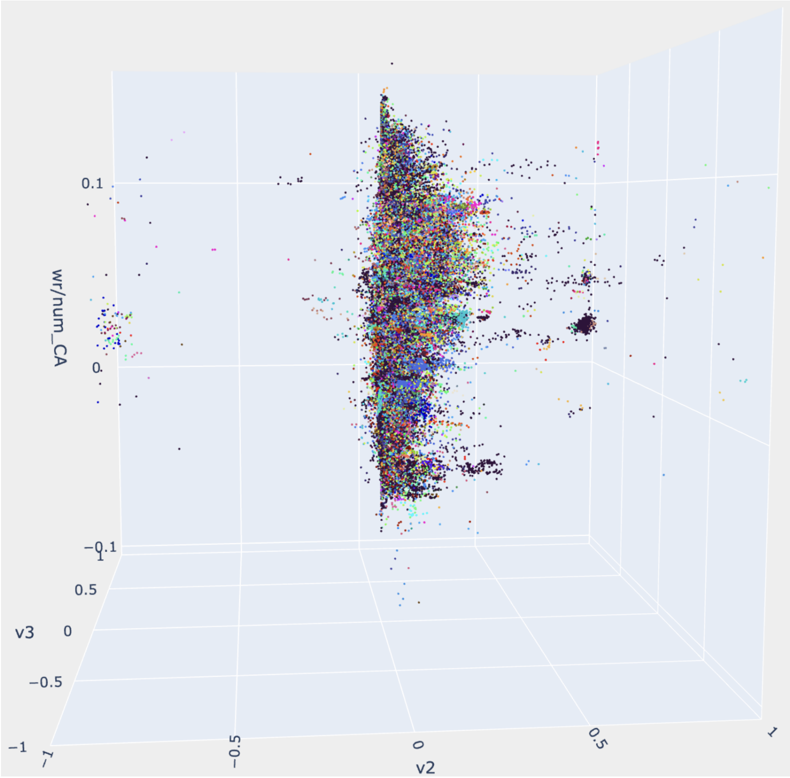


Figure 1: The entanglement-based conformational landscape of the Protein Data Bank (PDB). Each protein corresponds to a point in 3-space with axes Writhe normalized by its length, second and third Vasiliev measure. The proteins are colored according to the species they represent. In black are shown human proteins.

### 2 The second and third Vassiliev measures of knotoids and open curves in 3-space

In the case of knots, the second and third Vassiliev invariants are known to be correlated when comparing knots of the same crossing number (that is the minimum number of crossings over all embeddings and all projections of a knot). The correlation is manifested as a pattern of the $v_2, v_3$ values of knots of fixed crossing number in the $v_2, v_3$ plane that is called Willerton's fish, see [1]. Therein, it is also shown that the body contains amphichiral knots and the tail torus knots. It is an open question to determine if and to what extent this known correlation between $v_2, v_3$ in the case of knots also manifests in the $v_2, v_3$ values in the case of proteins. Note that proteins are modeled by simple open curves in 3-space and therefore are not mathematical knots (the latter are simple closed curves in 3-space). Moreover, the $v_2, v_3$ plane of the entanglement landscape of proteins does not depict proteins grouped by any topological consideration, such as crossing number or some analogue for open curves.

We use numerical experiments to examine the extent to which the pattern observed for proteins in the $v_2, v_3$ plane is related to the underlying mathematical correlation of $v_2, v_3$ in the case of knots. First, we examine the pattern correlation of the $v_2, v_3$ plane of knots when not restricting the crossing number of knots. Superimposing the $v_2, v_3$ values for prime knots of crossing number up to 12 we still observe a pattern similar to Willerton's fish (see Figure 2). Note that the points in the patters are all on an integer lattice, as expected for knots. The preservation of the pattern is expected, due to the proven inequalities relating $v_2, v_3$ of knots and their crossing number in [1].

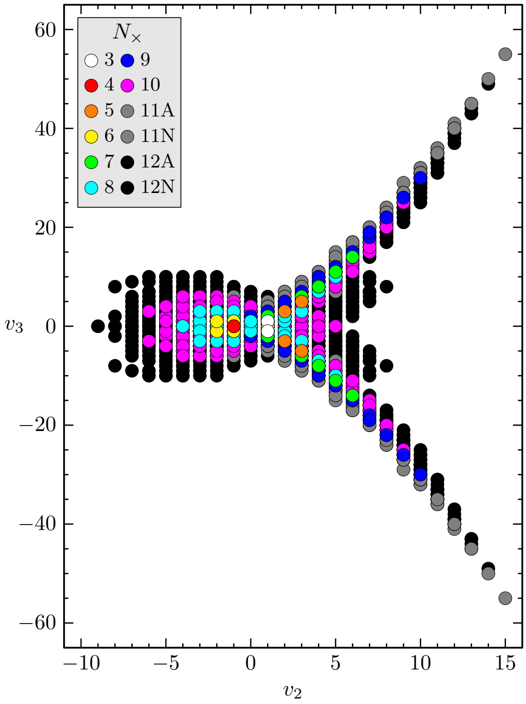


Figure 2: The $v_2, v_3$ values of all prime knots with crossing number less than or equal to 12. We observe that even when superimposing the results of different crossing number, a Willerton's fish pattern appears as in [1].

Second, we examine the pattern correlation of the $v_2, v_3$ plane of knotoids (equivalence classes of open arc diagrams [2]) of random crossing number. For knotoids, the $(v_2, v_3)$ values are in a lattice $1/2\mathbb{Z} \times 1/2\mathbb{Z}$. We generate equilateral random walks of length 10 to 100 and project them in a plane to obtain random polygonal knotoids. From the $v_2, v_3$ values of the random diagrams of knotoids we observe a pattern similar to that of knots arise (see Figure 3 (a). This can be explained as the inequaltities in [1] hold also for knotoids grouped by their crossing number (for knotoids it is defined as the minimum number of crossings of any diagram of a knotoid class). With similar arguments as for knots in [1], amphichiral knotoids will be in the body of the fish, while torus knotoids will be in the tails. We observe that the fish pattern of knotoids is not as well defined as that of knots shown in Figure 2. This is explained, at least in part, by the fact that Figure 3 (a) shows superimposed knotoids of random crossing numbers and not that of all knotoids of any given range of crossing numbers.

Lastly, we examine the pattern correlation of the $v_2, v_3$ plane of open curves in 3-space (this is the relevant case to proteins). For open curves in 3-space, $v_2, v_3$ are real numbers. By generating equilateral random walks of length 10 to 100 we obtain their $v_2, v_3$ values, shown in Figure 3 (b). We see that a pattern similar to Willerton's fish appears. This can be explained to an extent by the results for knotoids, as the $v_2, v_3$ values of open curves in 3-space are obtained by averaging that of knotoids over all possible projection directions [3]. Comparing this to Figure 1 in the main text, we notice overall the same pattern. Interestingly, some differences are evident which may indicate biological signatures in the correlation. More precisely, we see that the protein landscape (Figure 1 in main text) exhibits a non-symmetry of the two arms, with a bias towards positive $v_3$ values, as well as several flares and islands around the body of the fish, which is not obvious in Figure 3 (b). This suggests that there may be aspects of the pattern of the $v_2, v_3$ plane related to proteins in particular, as opposed to correlations of $v_2, v_3$ for random curves in 3-space and could be of interest for further study.

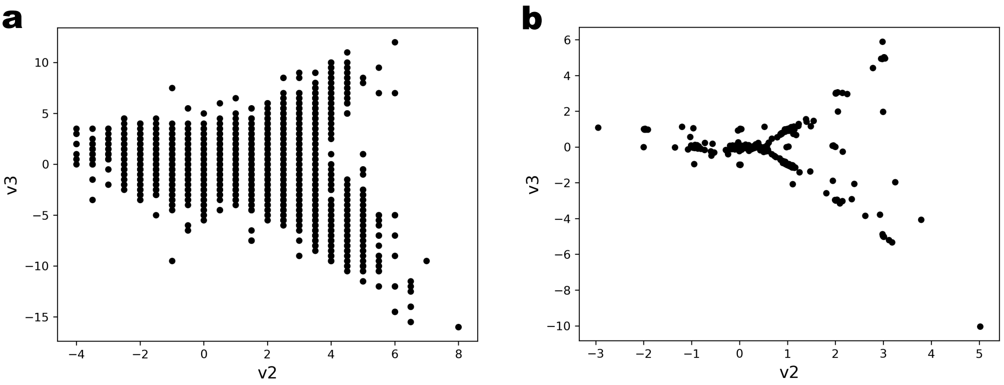


Figure 3: (a) The $(v_2, v_3)$ values of random knotoids (diagrammatic equivalence classes) of 10–100 edges. (b) The $(v_2, v_3)$ values of equilateral random walks (openn curves in 3-space) of length $N = 10$–100 edges.

### 3 CATH classification prediction

The data set was split 80/20 (stratified) into training and testing sets. A multi-layer perceptron is used with two hidden layers of 100 and 50 units and ReLu activations, trained with Adam optimizer and L2 regularization ($\alpha = 0.001$) with an adaptive learning rate. Hyperparameters were selected by 3-fold cross-validated grid search on the training set. To check overfitting, a separate run with the same hyperparameters was trained on a 90/10 train and validation split of the training data, with log-loss reported per epoch (see Figures 4,5,6,7,8,9).

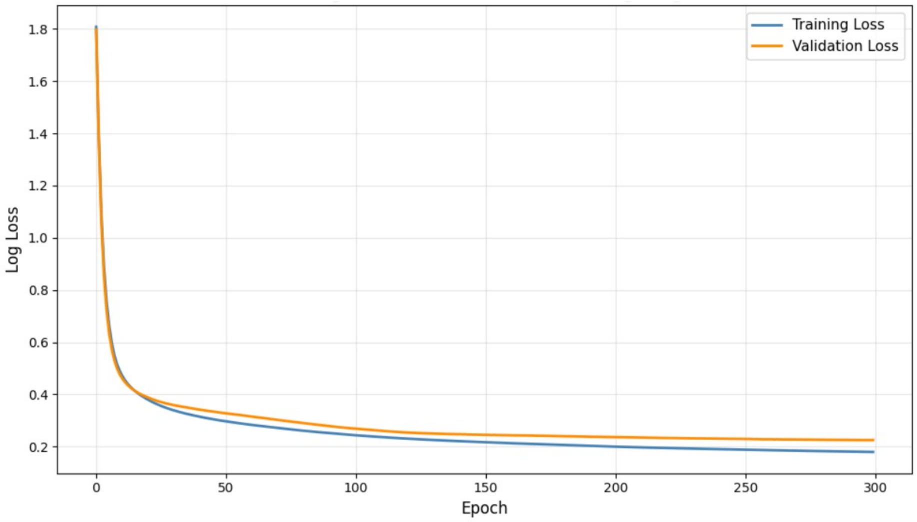


Figure 4: Loss Curves for $\alpha$ Class. Over 300 epochs, training and validation loss fell in parallel to 0.180 and 0.225 with a small gap of 0.04 indicating that model is trained cleanly without overfitting.

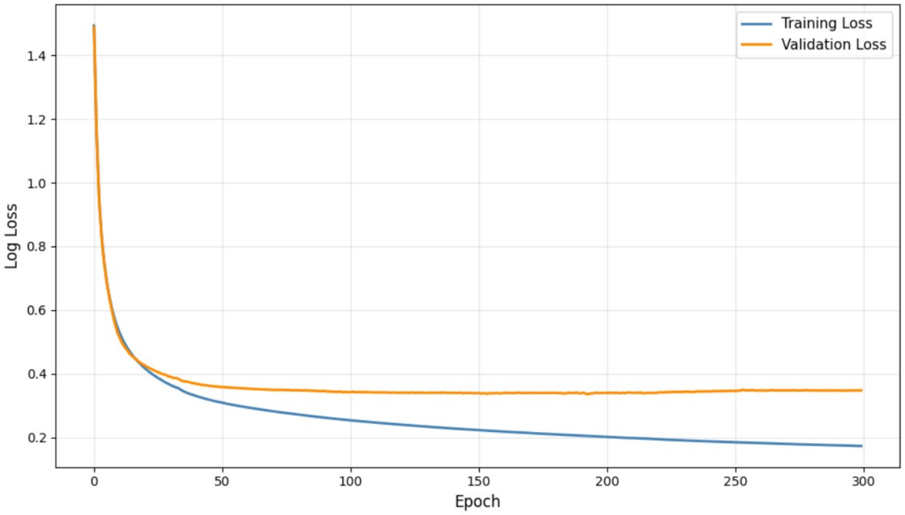


Figure 5: Loss Curves for $\beta$ Class. Training loss fell to 0.173 while validation plateaued at 0.348 after epoch 40, with a gap of 0.175. The train-test accuracy is gap is less than 3% showing that the model generalizes well.

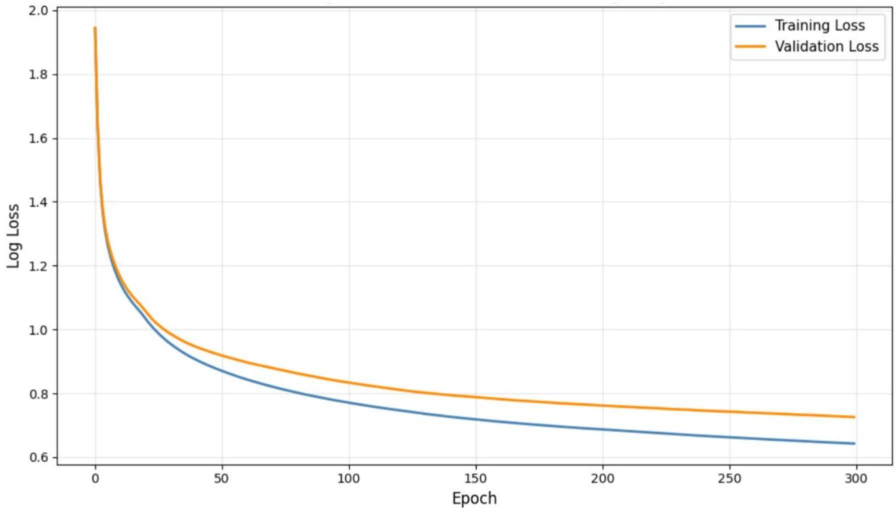


Figure 6: Loss Curves for $\alpha$-$\beta$ Class. Both training and validation curves decrease slowly and remain close (final gap = 0.083), suggesting that the model is still learning at epoch 300 with no overfitting.

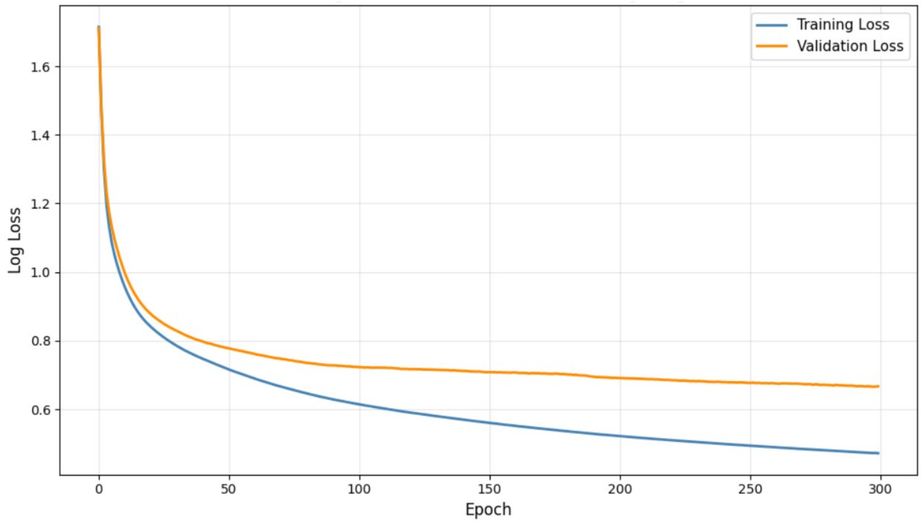


Figure 7: Loss Curves for $\alpha$-$\beta$ Class 3.40. Training loss continues to decrease while validation plateaus around epoch 100 with a final gap of 0.1949. But the train-test gap is less than 2% which suggests that the model generalizes well.

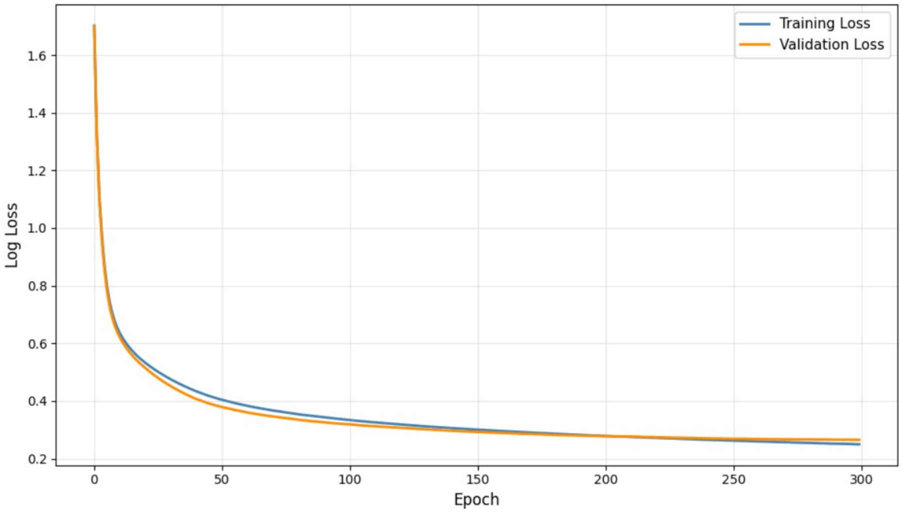


Figure 8: Loss Curves for $\alpha$-$\beta$ Class excluding 3.40. Training and validation curves decrease smoothly and remain tightly aligned (gap = 0.0156).

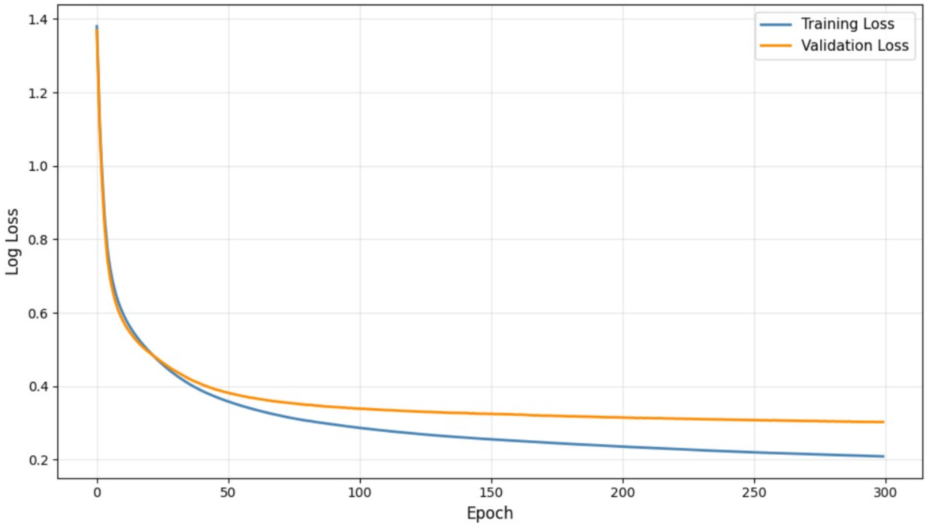


Figure 9: Loss Curves for $\alpha$-$\beta$ Class excluding 3.40.50. Training loss decreases slowly while validation loss flattens around epoch 150 (final gap = 0.0937), showing stable generalization.

Our ML model based on writhe, $v_2, v_3$ and length can predict some classes better than others. Namely, for $\alpha$ proteins we find that among the 10 subclasses, classes 1.10.630.10, 1.10.565.10, and 1.10.530.10 are the best-performing classes, with F1 scores of 0.997, 0.982, and 0.930 respectively. In contrast, 1.10.10.10 (F1 score: 0.683) performs the worst as lower precision and recall suggest greater confusion with other classes. For $\beta$ proteins we find that classes 2.40.70.10, 2.40.10.10, and 2.60.40.10 show the strongest performance, with consistently high F1 scores of 0.968, 0.932, and 0.916 respectively having a very few misclassifications. However, 2.40.155.10 has the weakest performance with F1 score of 0.824, mainly due to lower recall, meaning that many true samples are missed. For $\alpha - \beta$ proteins, we find that classes 3.10.200.10, 3.40.710.10 and 3.40.190.10 perform well in particular, with F1 scores of 0.997, 0.920, and 0.825 respectively. For $\alpha - \beta$ proteins without 3.40 (CATH): Classes 3.10.200.10 (1.000) and 3.40.710.10 (0.968) are the strongest performing classes with almost perfect F1 scores while 3.20.20.80is the weakest with F1 score of 0.784.
